# Alternative 3′ UTRs contributes to post-transcriptional gene expression regulation under high salt stress

**DOI:** 10.1101/2022.03.04.482946

**Authors:** Taotao Wang, Wenbin Ye, Jiaxiang Zhang, Han Li, Weike Zeng, Sheng Zhu, Guoli Ji, Xiaohui Wu, Liuyin Ma

## Abstract

High salt stress continually challenges growth and survival of many plants, but the underlying molecular basis is not fully explored. Alternative polyadenylation (APA) produces mRNAs with different 3′UTRs (alternative 3′UTRs) to regulate gene expression at post-transcriptional level. However, the roles of such process in response to salt stress remain elusive. Here, we reported that alternative 3′UTRs responded to high salt stress in the halophyte-*Spartina alterniflora*, which tolerant to hash salt environment. High salt stress induced the global APA and increased the prevalence of APA events. Strikingly, high salt stress significantly led to 3′ UTR lengthening of 207 transcripts through increasing the usage of distal poly(A) sites. Transcripts with alternative 3′ UTRs were mainly enriched in salt stress related ion transporters. Alternative 3′ UTRs of *SaHKT1* increased RNA stability and protein synthesis *in vivo*. Regulatory AU-rich elements were identified in the alternative 3′ UTRs and alternative 3′ UTRs increased protein level of *SaHKT1* in an AU-rich element dependent manner. Finally, 3′ UTR lengthening might result from variations in poly(A) signals and poly(A) factors. Overall, these results suggest that APA is a potential novel high salt stress responsive mechanism by modulating mRNA 3′ UTR length. These results also reveal complex regulator roles of alternative 3′ UTRs coupling alternative polyadenylation and regulatory elements at post-transcriptional level in plants.

**One sentence summary:** Alternative 3′ UTRs acts as a potential novel mechanism in gene expression regulation of high salt tolerant genes

## INTRODUCTION

Plants are sessile, and their growth and development are highly dependent on the surrounding environment (Munns and Tester, 2008; Deinlein et al., 2014). Soil salinization is one major environmental pressures restricting agricultural production. High salt stress causes hyperionic and hyperosmotic stresses, which induces cell membrane disintegration, produces reactive oxygen species, weakens metabolic processes, inhibits photosynthesis and reduces nutrient absorption, thereby delaying crop growth and reducing yield (Zhu, 2002; Ding et al., 2014). Gene expression of these processes is highly regulated by transcription factors at transcription level (Zhu, 2002). However, little is known about high salt stress associated gene expression regulation at the post-transcriptional level.

The 3′ untranslated regions (3′ UTRs) are important component of mRNAs in bacteria, archaea, and eukaryotes (Mayr, 2017). 3′ UTRs play essential regulatory roles in determining the diverse fate of mRNA at post-transcriptinal level by modulating mRNA stability, translation and localization (Tian and Manley, 2017; Mayr, 2019). The function of 3′ UTR on gene regulation is largely dependent on *cis*-acting elements (e.g. AU-rich elements and miRNA targeting sites) inside 3′ UTR and their associated *trans*-acting factors (RNA-binding proteins, RBPs) (Mayr, 2016, 2017). The presence and accessibility of these regulatory elements is controlled by alternative polyadenylation (APA) (Mayr, 2017).

Polyadenylation of pre-mRNA is a critical step in the maturation of eukaryotic mRNA and determines the position of mRNA 3′-end (Xing and Li, 2011; Hunt, 2014; Deng and Cao, 2017; Mayr, 2019). Moreover, up to 70% of eukaryotic genes contain multiple poly(A) sites and undergo APA (Wu et al., 2011; Fu et al., 2016). APA generates mRNA with different 3′ UTRs and many highly conserved *cis-acting* elements such as microRNA targeting sites and AU-rich elements, widely occur in the alternative 3′ UTRs in animals (Mayr and Bartel, 2009; Mayr, 2017; Tian and Manley, 2017; Mayr, 2019). For example, in the case of oncogenesis, the proximal poly(A) site is preferred to regulate 3′ UTR shortening of the Cyclin mRNA, which results in the loss of 3′ UTRs miRNA binding sites inside the alternative 3′ UTRs. This leads to the release of miRNAs mediated post-transcriptional repression of Cyclin expression and thereby resulted in uncontrolled cell division (Mayr and Bartel, 2009). In animal, APA events that shorten 3′ UTRs are associated with proliferating T cells, pluripotent stem cells, and cancer cells (Sandberg et al., 2008; Ji and Tian, 2009; Mayr and Bartel, 2009). 3′ UTR lengthening occurs in embryonic development, the nervous system, and in cellular senescence (Ji et al., 2009; Hilgers et al., 2012; Chen et al., 2018). Therefore, alternative 3′ UTRs have crucial regulatory roles in diversifying the fate of mRNAs and proteins, including effects on mRNA degradation, translation efficiency, and localization (Mayr, 2017). The occurrence of APA mediated alternative 3′ UTRs is regulated by *trans*-acting poly(A) factors such as CFIm25, CFIm68, CPSF100, and CPSF30; mutations in these factors result in significant 3′ UTR shortening or lengthening of mRNAs (Masamha et al., 2014; Xia et al., 2014; Gennarino et al., 2015; Lin et al., 2017). However, how alternative 3′ UTRs regulate gene expression is largely unknown in plants.

The study of plant APA is currently mainly focused on gene expression regulation in non-canonical polyadenylation or shifting between non-canonical and 3′ UTRs polyadenylation. For example, APA and epigenetic modifications complex competes to bind a transposon element in the long intron region of IBM1 mRNA, and thus regulates the expression of short-segment mRNA via intronic APA or full-length mRNA resulted from shielding the intronic APA sites by epigenetic modification (Lei et al., 2014; Ma et al., 2014; Duan et al., 2017; Zhang et al., 2021). In the flowering autonomous pathway that consisted with FCA-FPA-FLC, FCA and FPA are RNA-binding proteins that promote proximal polyadenylation of FLC antisense RNA to reduce FLC expression and induce flowering (Deng and Cao, 2017). In plants, hypoxic stress increases the usage of non-canonical poly(A) sites in genic regions other than the 3′ UTR, thereby affecting the stability of the affected transcripts (de Lorenzo et al., 2017). In *Arabidopsis thaliana*, dehydration stress results in significant 3′ UTR lengthening and these alternative 3′ UTRs act as non-coding RNAs to regulate the expression of downstream genes (Sun et al., 2017). However, questions regarding the relationship between alternative 3′ UTRs and salt stress remain unresolved.

The invasive monocotyledonous halophyte - *Spartina alterniflora* (*Spartina*) is a good model to study high salt tolerance mechanism, as it could tolerant to harsh salinity environment in salt marsh (Bedre et al., 2016; Ye et al., 2020). In this study, we used PacBio SMRT sequencing and poly(A) tag sequencing (PAT-seq) to get insight into how APA responds to high salt stress in *Spartina*. Moreover, the important contributions of APA in high salt stress are systematically deciphered from both genome-wide and gene-specific aspects. Specifically, the effects of alternative 3′ UTRs on mRNA accumulation, RNA stability, and protein synthesis were analyzed and the biogenesis of alternative 3′ UTRs were also unveiled.

## RESULTS

### Profile of poly(A) sites under high salt stress

An overview of the experimental procedure was illustrated in Figure 1. Briefly, total RNAs were isolated from 24 hr salinity stress treated *Spartina* seedling (0, 350 mM, 500 mM and 800 mM) to simulate the control, lower, medium, and higher salt stress as described in previous study (Ye et al., 2020). Twelve poly(A) tag libraries (PAT-seq) were constructed using total RNAs from various salt stress in *Spartina* and sequenced on illumina HiSeq 2500 platform (Figure 1). A total of 38,876,361 poly(A) reads were obtained by mapping PAT-seq reads to *Spartina* full-length transcripts from PacBio sequences (**Table S1**). Adjacent poly(A) sites within 24 nt were grouped into poly(A) site clusters (PACs) to reduce the effects of microheterogeneity (Wu et al., 2011), and 47,176 PACs supported by more than 10 reads were identified. Over 85% of 3′ UTR-PACs were localized in the [0, 10 bp] window at the 3′ terminal of PacBio full-length non-chimeric (FLNC) transcripts (**Figure S1**), indicating the strategy for analyzing the 3′ end transcriptome by combining the PacBio and PAT-seq is effective. The single-nucleotide profile around poly(A) sites resembles the general profile of other plant species such as rice or Arabidopsis (Wu et al., 2011; Fu et al., 2016), where a conserved U-rich peak upstream of the poly(A) site as well as the dinucleotide YA (Y=C or U) at the cleavage site were observed (Figure 2a). Moreover, the proportion of canonical AAUAAA and its 1-nt variants are also comparable to that of rice (Figure 2b).

**Figure 1.**
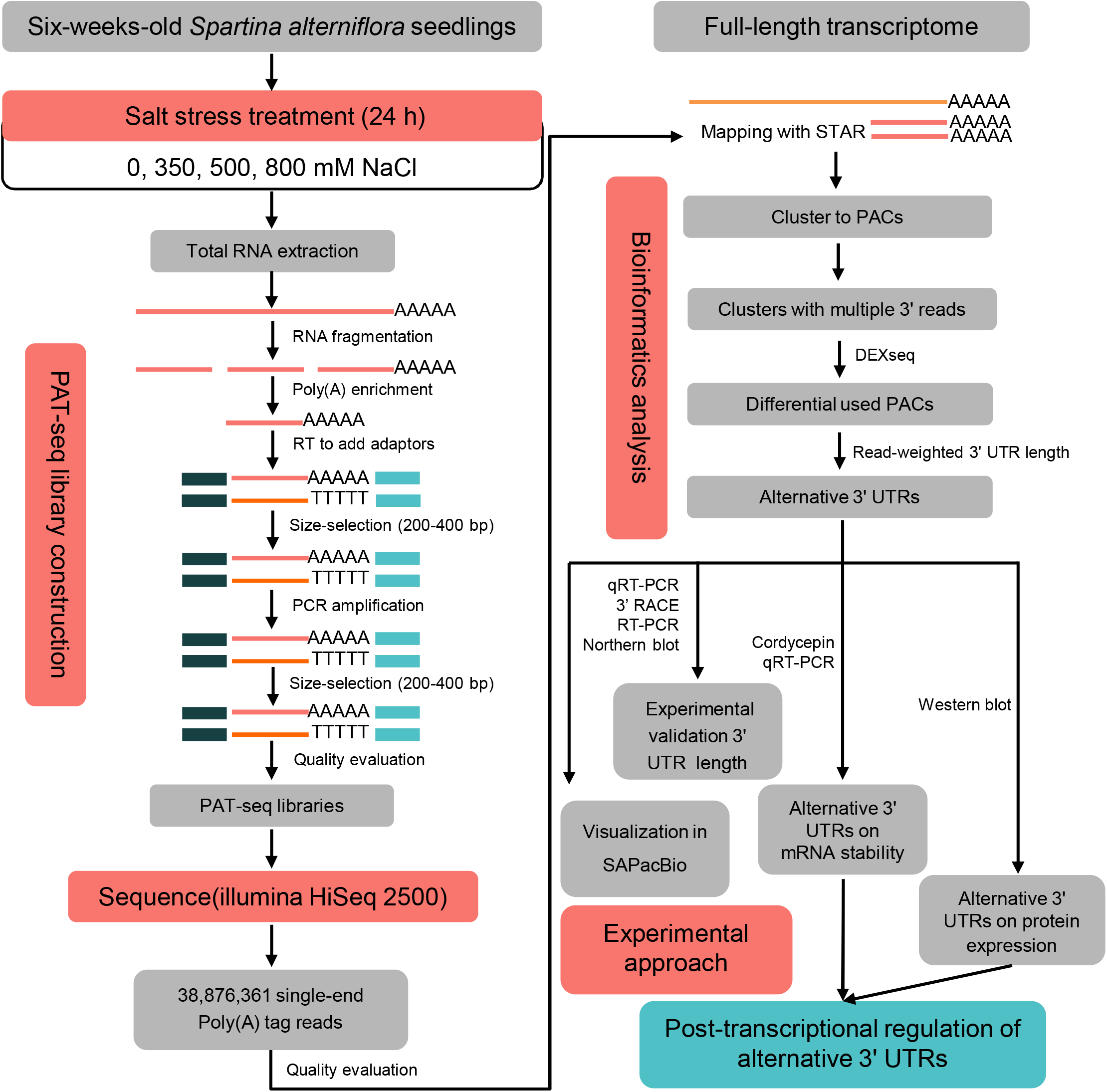
Overview of the experimental and bioinformatics workflow. Major procedures in this study include: (1) salt stress treatment; (2) PAT-seq libraries construction; (3) bioinformatics analysis; (4) experimental approaches such as experimental validation, RNA stability assay, Western blot, and PAT-seq: poly(A) tag sequencing that specific containing the 3′ end information.

**Figure 2.**
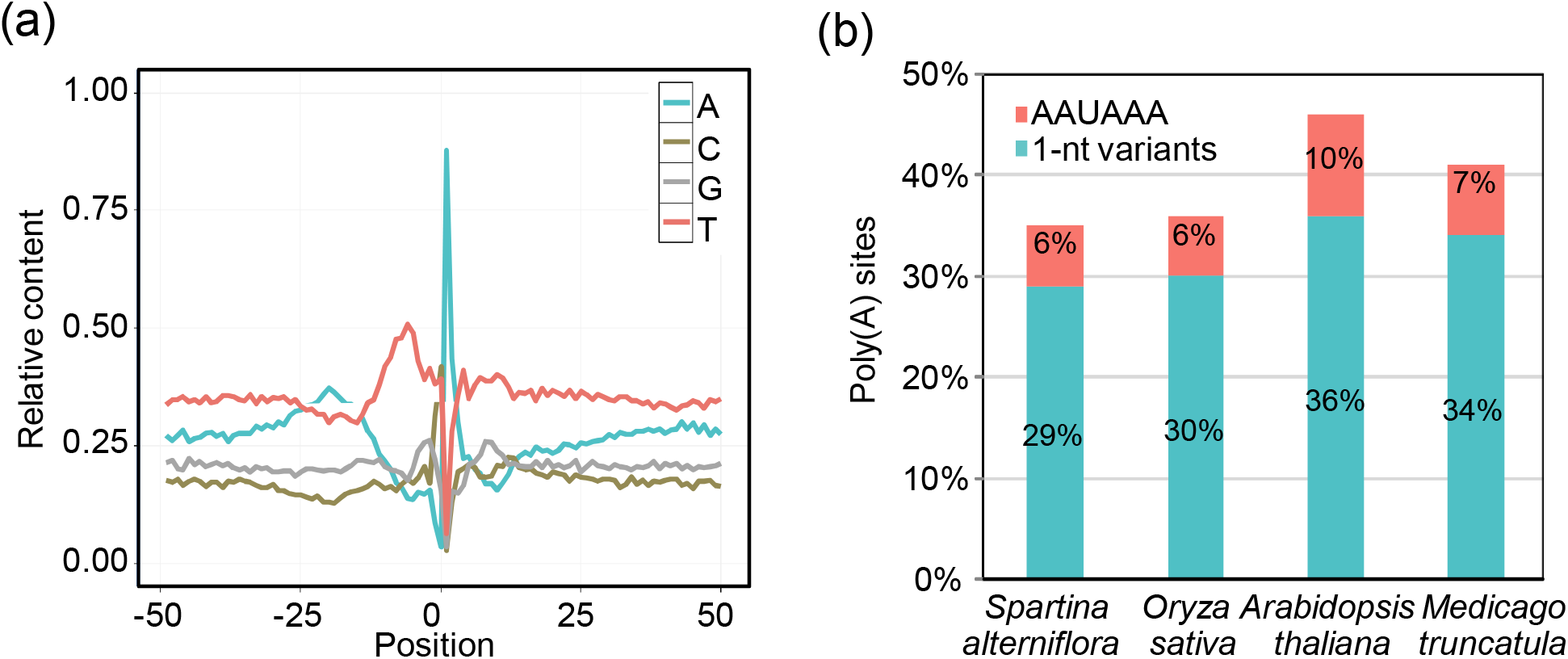
Characterization of poly(A) signal and sequence distribution pattern. (a) Single nucleotide profile of the sequences surrounding poly(A) sites. Y-axis denotes the fractional nucleotide content at each position. On the x-axis, ‘0’ denotes the poly(A) site, ‘-’ denotes upstream. (b) Distribution of canonical AAUAAA and its 1-nt variants in near upstream of PACs among different plants.

### APA is widespread under high salt stress

APA is prevalent in *Spartina* with 53.86% (9248/17168) of expressed unigenes possessing more than one PAC. To further understand the extent of APA involved in response to salt stress, DEXseq (Anders et al., 2012) was introduced to identify differential expressed PACs (DEPACs) from APA-containing unigenes under salt stress. A total of 8733 DEPACs corresponding to ∼42% (3886/9248) of APA-containing unigenes were found in response to salt stress (**Table S2**). Importantly, the number of DEPACs increased along salt gradients (Figure 3a), suggesting salt stress induced APA events. High salt stress affects the differential expression of 7339 PACs, while with a slightly more up-regulated PACs than down-regulated ones(3766 vs 3573, Figure 3b).

**Figure 3.**
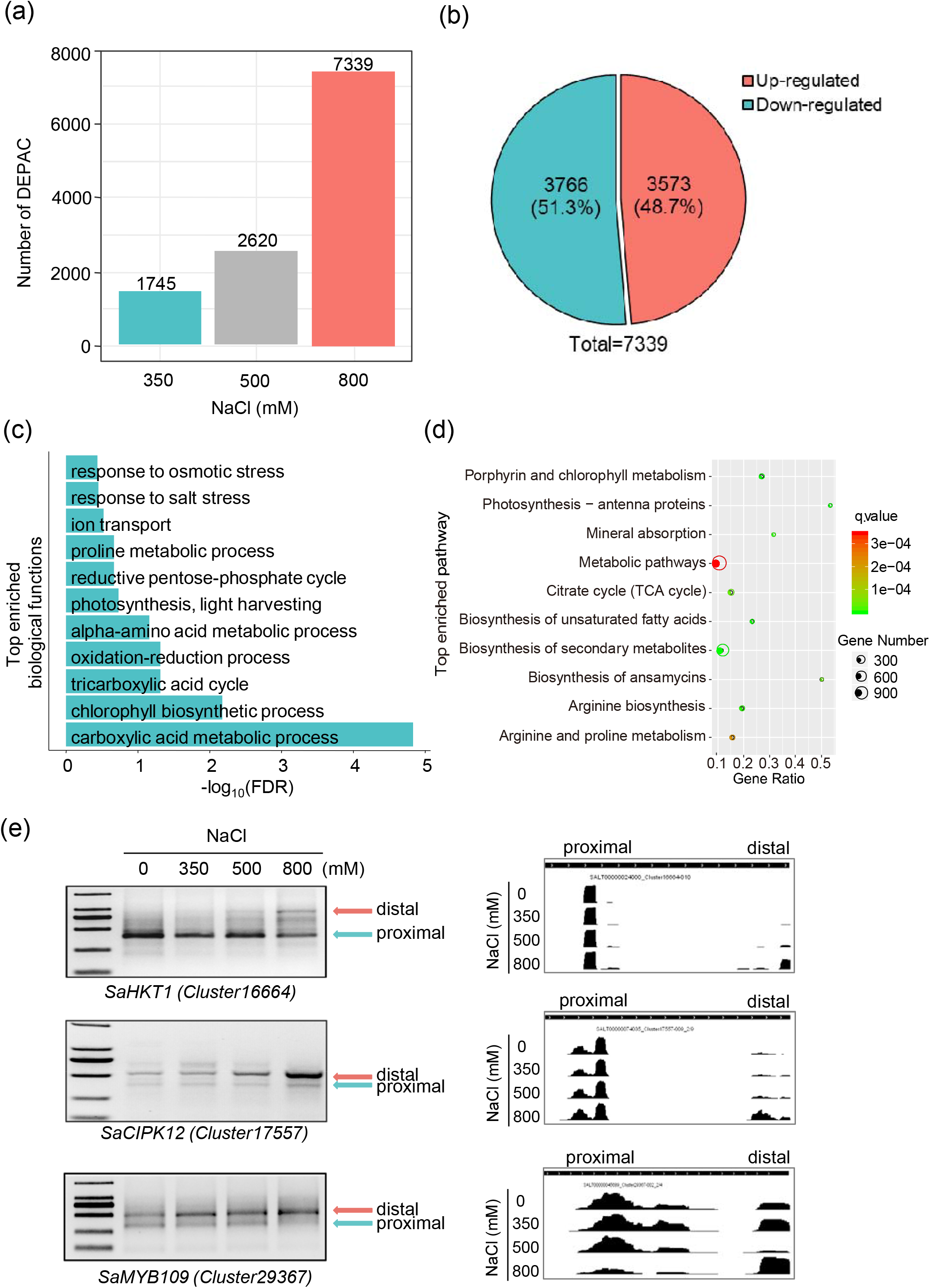
Differential expressed PACs in response to high salt stress (a) Number of differentially expressed PACs (DEPACs) under different salt stress treatments. (b) Number of up-regulated or down-regulated DEPACs under high salt stress. (c) Gene ontology analysis of high salt stress associated DEPACs unigenes (*P* < 0.05). (d) KEGG pathway analysis of high salt stress associated DEPACs unigenes (*P* < 0.05). (e) Left panel: Salt-related DEPACs were validated by 3′RACE. Right panel: Visualization of PAT-seq reads accumulation of distal poly(A) sites for salt-responsive genes under high salt stress. *SaHKT1: high-affinity potassium transporter 1*; *SaCIPK12: CBL-INTERACTING PROTEIN KINASE 12*; distal: distal poly(A) sites; proximal: proximal poly(A) sites.

Gene ontology (GO) and KEGG analysis reveals that PACs associated with high salt stress are overrepresented in the oxidative-reduction process, amino acid metabolic processes, ion transport, responses to osmotic stress, responses to salt stress, the TCA metabolic process, and photosynthesis (Figure 3c-d). These pathways have been found in responses to salt stress (Munns and Tester, 2008; Deinlein et al., 2014; Soni et al., 2015).

We validated APA of three genes which have been reported to play vital roles in response to salt stress (Rus et al., 2001; Xiang et al., 2007; Harb, 2010): the ion transport encoding gene *SaHKT1 (Cluster1664-010)*, the kinase *SaCIPK12 (Cluster17557-009),* and the transcription factor *SaMYB109 (Cluster29367-002)*. Both the results from 3′ RACE experiments and visualization of PAT-seq reads showed that high salt stress induces usage of distal poly(A) sites (Figure 3e). Overall, our results reveal that APA responds to high salt stress by affecting the usage of poly(A) sites in Spartina.

### High salt stress induced 3′ UTR lengthening in *Spartina*

To assess whether APA responds to high salt stress, we first adopted a specific assay from a previous study (Thomas et al., 2012) to evaluate differences in poly(A) site choice between salt treatments and the control. Each gene was assigned a value between 0 and 1, which indicates the difference in poly(A) site choice between two conditions. The running sum of the number of genes with values falling within a given interval was plotted to reflect the global difference of poly(A) site usage between two conditions. Global differences in the use of APA between the higher salt stress and the control were much greater than between the lower salt treatment and the control (Figure 4a). The difference between the 800 mM NaCl treatment and the control seemed to be much more dramatic than that between the control and the 500 mM NaCl or 350 mM NaCl treatments (Wilcoxon test, *P* = 4.09e-236; Figure 4a), suggesting that high salt stress significantly induced widespread dynamics of poly(A) site choice.

**Figure 4.**
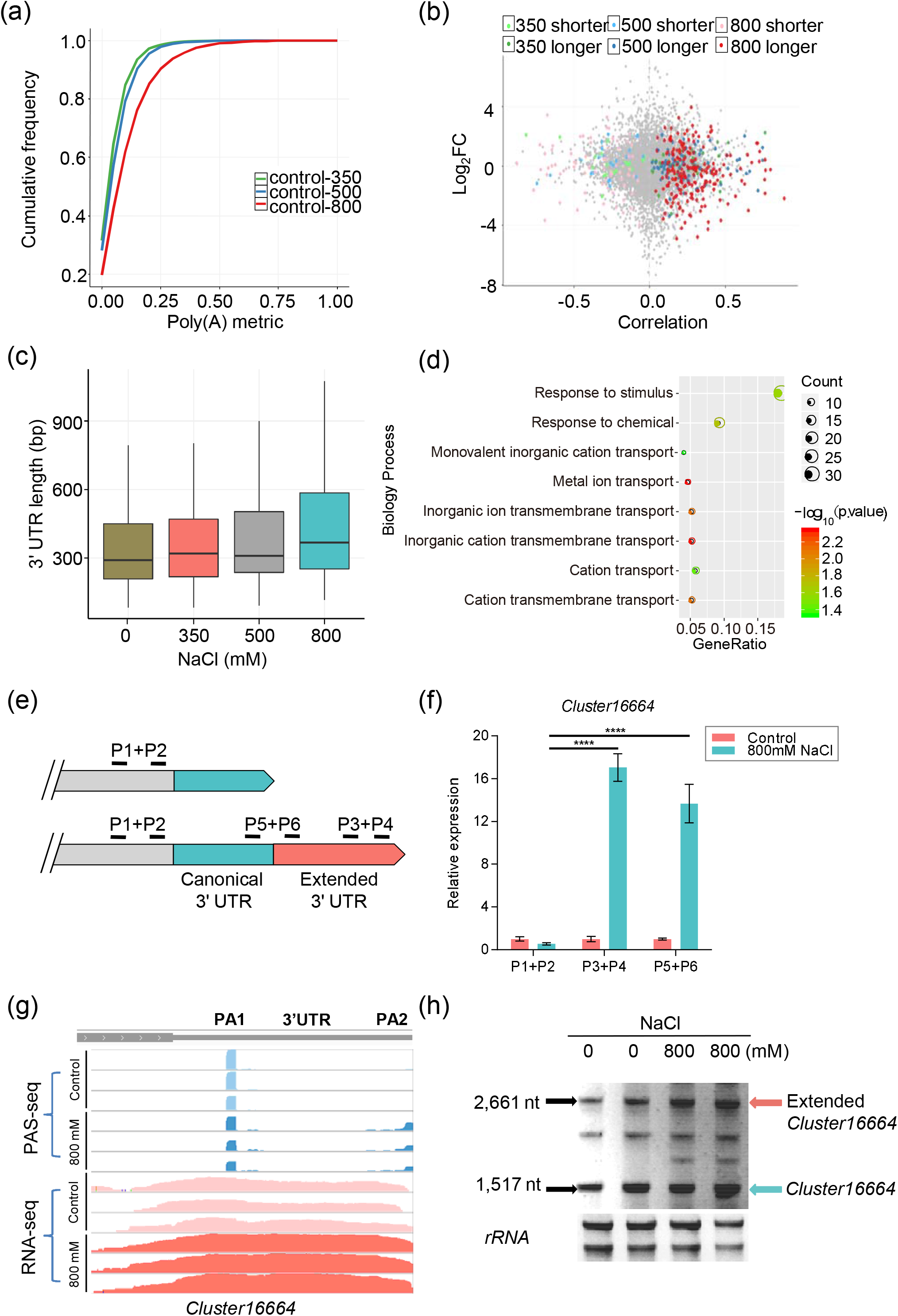
High salt stress induced 3′ UTR lengthening. (a) Plot of the running sum of genes with increasing differences in the poly(A) site usage between conditions. (b) Significant 3′ UTR lengthening and shortening between each salt treatment and the control. Pearson product–moment correlation coefficient is plotted against the log_2_ fold change between conditions. Genes with significant switching to longer or shorter 3′ UTRs are colored. (c) Distribution of 3′ UTR lengths for unigenes with significant lengthening in 800 mM NaCl treatment across different conditions. (d) Gene ontology analysis of high salt stress associated 3′ UTR lengthening unigenes. (e) Schematic showing the experimental strategy to validate 3′ UTR lengthening with three pairs of primers: the P1+P2 primer pair detects the expression of both cUTRs (canonical 3′ UTR) and aUTRs (alternative 3′ UTR); the P3+P4 pair detects the expression of aUTRs only; the P5+P6 pair detects expression of aUTRs and verifies the aUTR result from the same transcripts. (f) 3′ UTR lengthening under high salt stress was validated by qRT-PCR with P3+P4 primers. P5+P6 primers was used to validate that 3′ UTR lengthening resulted from the same transcript but not independent transcripts. (g) Visualization of RNA-seq reads accumulation in alternative 3′ UTR of *SaHKT1* transcripts under high salt stress. (h) Northern blot assay detecting the mRNA level of *SaHKT1* transcripts under high salt stress. Cluster16664 represents *SaHKT1* transcripts with both aUTR + cUTR; extended Cluster16664 represents *SaHKT1* transcripts with only aUTR.

Next, we examined 3′ UTR variation under different salt stress conditions. Among the 8733 transcripts with differential APA usage obtained from *Spartina* under different salt stress and control conditions, 896 showed significant 3′ UTR lengthening or shortening (Figure 4b). Importantly, salt stress induced more mRNA 3′ UTR lengthening events than 3′ UTR shortening events, and the number of varied 3′ UTRs increased along the salt stress gradient (Figure 4b and **S2**). In particular, 457 unigenes displayed significant 3′ UTR lengthening under salt stress (FDR < 0.05), and 207 of them were associated with high salt stress (**Table S3**).

### 3′ UTR lengthening of ion transporter transcripts under high salt stress

To understand the extent of 3′ UTR lengthening, the average 3′ UTR length weighted by expression level under each salt stress condition was measured (Ulitsky et al., 2012). We found that high salt stress leads to 3′ UTR lengthening for more than 60 bp (Figure 4c). Interestingly, gene ontology analysis showed that metal ion transport is significantly enriched in high salt stress-associated 3′ UTR lengthening unigenes (*P* = 4.51e-03; Figure 4d). These results indicated a potential link between 3′ UTR lengthening and ion transport mediated high salt tolerance.

In plants, it has been reported that high salt stress induced accumulation of independent 3′ transcription products through suppression of XRN3 exoribonuclease activity (Kurihara et al., 2012; Krzyszton et al., 2018). To exclude such possibility, we designed a series of experiments to prove that high salt stress induced 3′ UTR lengthening is indeed real APA events but not from independent truncating transcripts. We first designed qRT-PCR experiments to validate high salt stress-associated 3′ UTR lengthening with three pairs of primers (Figure 4e). The ion transporter-*SaHKT1* (*Cluster16664*) were selected as an example for further analysis. The mRNA levels of 3′ UTR-lengthened regions (detected by P3+P4 pair primers) increased by over 10-fold by qRT-PCR under high salt stress (Figure 4f). Moreover, the increased expression of 3′ UTR regions flanking the proximal poly(A) sites of *SaHKT1* (detected by P5+P6 pair primers) by RT-PCR under high salt stress further supports that these 3′ UTRs lengthening is originated from the extension of 3′ UTRs (Figure 4f). The read coverage of extended 3′ UTR regions from RNA-seq data also supported that the 3′ UTR lengthening is the real APA event (Figure 4g). Finally, high salt stress induced 3′ UTR lengthening of *SaHKT1* transcripts was also proved by northern blot (Figure 4h), and the result further proved that these 3′ UTR lengthening events are real, rather than technical noise or mRNA decay fragments from independent transcripts. Overall, high salt stress leads to mRNA 3′ UTR lengthening by the preferential usage of distal poly(A) sites, suggesting a potential novel salt stress-responsive mechanism by modulating of the 3′ UTR length.

### 3′ UTR lengthening did not affect global RNA accumulation

In animals, variation of the 3′ UTR length is one of several regulatory features that regulate RNA stability, translation efficiency, and protein localization (Mayr, 2017; Tian and Manley, 2017). However, it is not clear how the varition 3′ UTR length affects mRNA and protein level in plants. First, we evaluated the effect of 3′ UTR lengthening on global mRNA accumulation, while we did not observe global RNA accumulation variation in the transcripts that undergo 3′ UTR lengthening (Figure 5a). We then selected four transporter genes that show 3′ UTR lengthening, *SaHKT1* (*Cluster16664*), *SaKT2* (*Cluster9684*), *SaZTP29* (*Cluster30331*), and *SaCIPK23* (*Cluster10136*) for detailed mRNA accumulation analysis. The mRNA level of both *SaKT2* and *SaCIPK23* increased under high salt stress, while that of *SaHKT1* and *SaZTP29* decreased (Figure 5b **and S3**). Again, these results indicate that 3′ UTR lengthening under high salt stress is not a global regulator for mRNA accumulation, but more likely occurs on a gene-by-gene basis.

**Figure 5.**
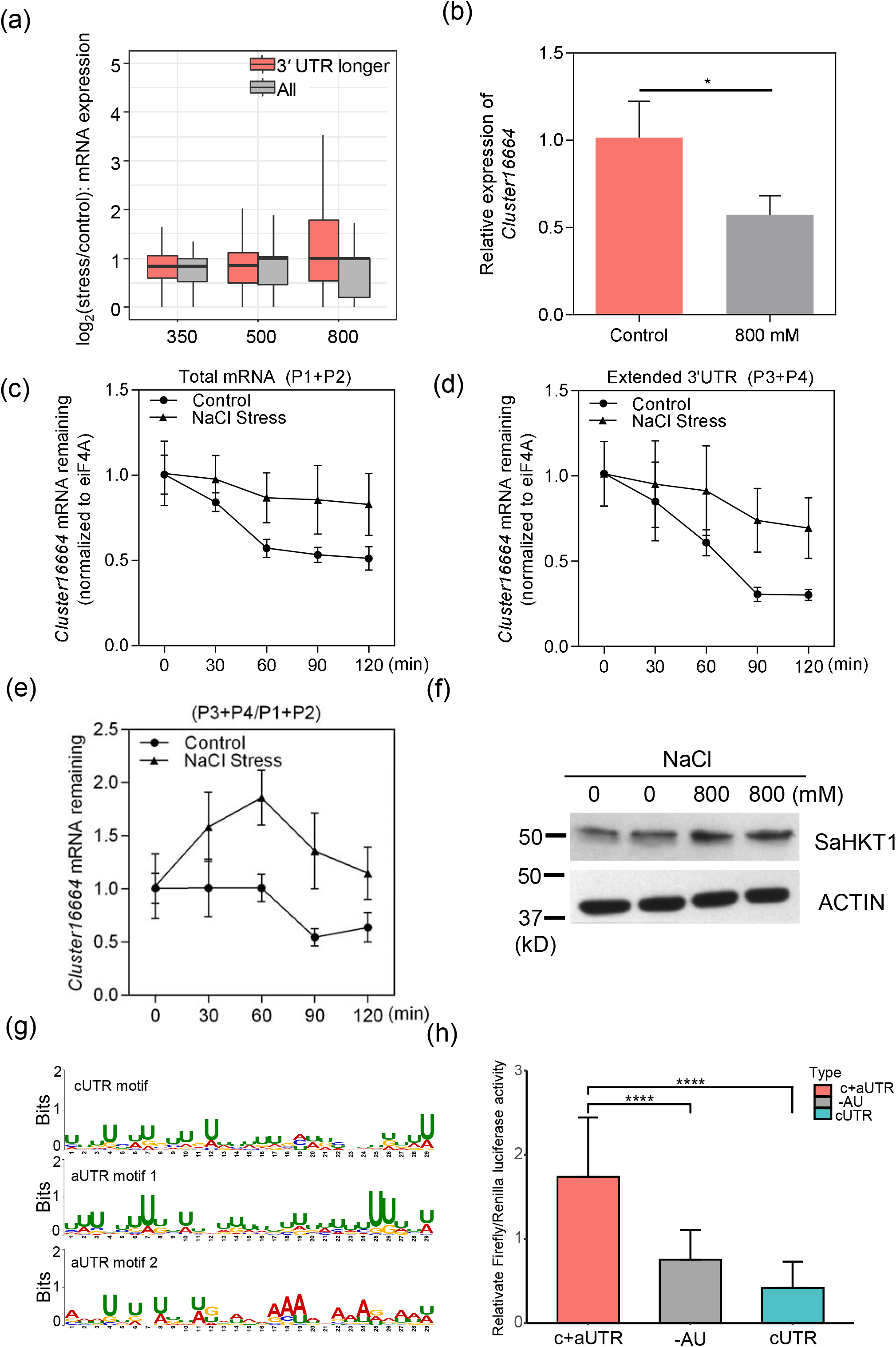
The effect of 3′ UTR lengthening. (a) Variation in the expression of genes with longer or shorter 3′ UTR. Expression value of each sample is calculated as the log_2_ (fold change) of gene expression value between the treated sample and the control. “All” denotes expression levels of all expressed genes without change of 3′ UTR in the respective sample, ‘*’ represents significant RNA level difference (*P* < 0.05). The x-axis represents different salinity conditions: 300: 300 mM NaCl treated seedlings etc. (b) Relative expression of *SaHKT1* under high salt stress detected by qRT-PCR. (c) The effect of high salt stress on RNA stability of remaining total transcripts (cUTR +aUTR) of *SaHKT1*. (d) The effect of high salt stress on RNA stability of remaining 3′ UTR transcripts (aUTR) of *SaHKT1*. (e) The effect of high salt stress on RNA stability of remaining transcripts ratio (P3+P4/P1+P2). (f) Western blot detecting the protein level of SaHKT1 under both control and high salt stress, ACTIN was used as the loading control, the gray value calculated with Gel-Pro Analyzer 4. (g) Motif analysis of canonical 3′ UTR and alternative 3′ UTR with MEME. (h) Dual luciferase assay of short (canonical, cUTR) 3′ UTRs, long (canonical + alternative, cUTR+aUTR) 3′ UTRs and 3′ UTRs with deleted AU-rich elements (-AU) of SaHKT1 in *Spartina* protoplast using PEG-Calcium-dependent transient expression assay. The ratio of Firefly/Renilla represents the effects of 3′ UTR length on protein expression. ‘***’ represents significant luciferase activity difference (*P* < 0.001).

### 3′ UTR lengthening increased mRNA stability

Alternative 3′ UTRs have been reported to affect RNA stability by inclusion or exclusion of regulatory elements (Mayr, 2019). We hypothesized that 3′ UTR lengthening of *Spartina* transcripts may also affect RNA stability under high salt stress. HKT1 are well-known high salt tolerant genes by reducing Na^+^ toxicity through K^+^ uptake and overexpression of HKT1;2 from the halophyte-*Eutrema parvula* conferred significantly high salt tolerance phenotype in Arabidopsis (Ali et al., 2018). To test our hypothesis, *SaHKT1* was selected to perform RNA stability assay *in vivo* using *Spartina* seedlings. We first evaluated the stability of total transcripts (cUTRs + aUTRs) under high salt stress using a pair of primers from exon regions (P1 + P2, Figure 5c). Generally, the results showed that total transcripts were more stable under high salt stress than the control (Figure 5c). Next, to test whether transcripts with extended 3′ UTRs contributed to the stability of total transcripts under high salt stress, a pair of primers (P3 + P4, Figure 5d) was designed to only amplify the transcripts with extended 3′ UTRs. Interestingly, the mRNA levels of *SaHKT1* were significantly reduced under high salt stress (Figure 5d). Conversely, the *SaHKT1* transcripts with extended 3’UTR were more stable under high salt stress than the control (Figure 5d). Moreover, the expression ratio of extended 3’UTR between P3+P4 and P1+P2 under high salt stress were consistently higher than that from control (Figure 5e), indicating that the extended 3’UTR of *SaHKT1* indeed increase RNA stability. Overall, these results indicate that 3’UTR extension may increase RNA stability under high salt stress, at least in the case of *SaHKT1*.

### 3′ UTR lengthening affects protein level

In mammalian systems, it has been reported that 3′ UTR shortening generally leads to higher protein level, while 3′ UTR lengthening may lead to the inclusion of negative regulatory elements, thus reducing protein levels (Mayr and Bartel, 2009). To test whether 3′ UTR lengthening can have effect on protein level under high salt stress in plants, we performed western blot assay of SaHKT1. Our results showed that the protein level of SaHKT1 was increased under high salt stress in *Spartina* (Figure 5f). Take together with the observation that 3′ UTR lengthening of *SaHKT1* transcripts under high salt stress, we rationally deduce that 3′ UTR lengthening increases the protein accumulation of SaHKT1. Overall, our results indicate that 3′ UTR lengthening may affect protein level.

### 3′ UTR lengthening increased protein synthesis via AU-rich element

In animal, regions of alternative 3′ UTRs (aUTR) contain important regulatory motifs (Mayr, 2017; Tian and Manley, 2017). We then scanned for potential motifs in alternative 3′ UTR regions associated with salt stress. Interestingly, two motifs, the U-rich element (29 nt, e-value = 2.7e-94) and the AU-rich element (29 nt, e-value = 2.3e-08), were identified in alternative 3′ UTR regions, while only the U-rich element (29 nt, e-value = 5.5e-15) was found in the canonical 3′ UTR (cUTR, Figure 5g), suggesting that AU-rich elements in alternative 3′ UTR regions may have important functions.

To further decipher whether alternative 3′ UTR regulates the RNA stability and protein synthesis in an AU-rich *cis*-element dependent manner, we established a protoplast transient expression system in *Spartina* and introduced a modified dual luciferase assay, which was used to study the interaction between miRNA and their 3′ UTR targeting sites previously (Liu et al., 2014), to assess the potential effect of 3′ UTR lengthening on protein synthesis. Briefly, different 3′ UTRs were cloned downstream from stop codon of firefly luciferase to calculate the luciferase activity variation and thereby evaluated the effects of alternative 3′ UTRs as well as associated regulatory elements on protein synthesis. To minimize the bias of different transformation efficiency, a renilla luciferase located in the same vector with firefly luciferase was used as an internal control. Similar to western blot experiments, we observed that extended 3′ UTRs of *SaHKT1(Cluster16664)* showed significant higher luciferase activity than the respective canonical 3′ UTRs (Wilcoxon test, *P* = 1.99e-07, Figure 5h). To further explore the roles of the AU-rich elements, we then deleted the 29 nt AU-rich element in alternative 3′ UTRs of *SaHKT1*. Interestingly, 3′ UTRs with the deletion of the element significantly reduced the luciferase activity (Wilcoxon test, *P* = 5.97e-05) than that without deletion (Figure 5h); suggesting that alternative 3′ UTRs of *SaHKT1* regulate protein synthesis in an AU-rich element dependent manner.

### The biogenesis of 3′ UTR lengthening

The choice of poly(A) site depends largely on the conservation of *cis*-acting poly(A) signals (Loke et al., 2005; Di Giammartino et al., 2011; Tian and Manley, 2017) and expression of *trans*-acting poly(A) factors (Mayr, 2017; Tian and Manley, 2017). The single nucleotide profile surrounding poly(A) sites showed that poly(A) signal of distal poly(A) sites was different from the proximal ones (Figure 6a-b). Compared to proximal sites, a higher A but lower U base composition was observed upstream (–35 nt to –10 nt) of distal poly(A) sites (Figure 6c).

**Figure 6.**
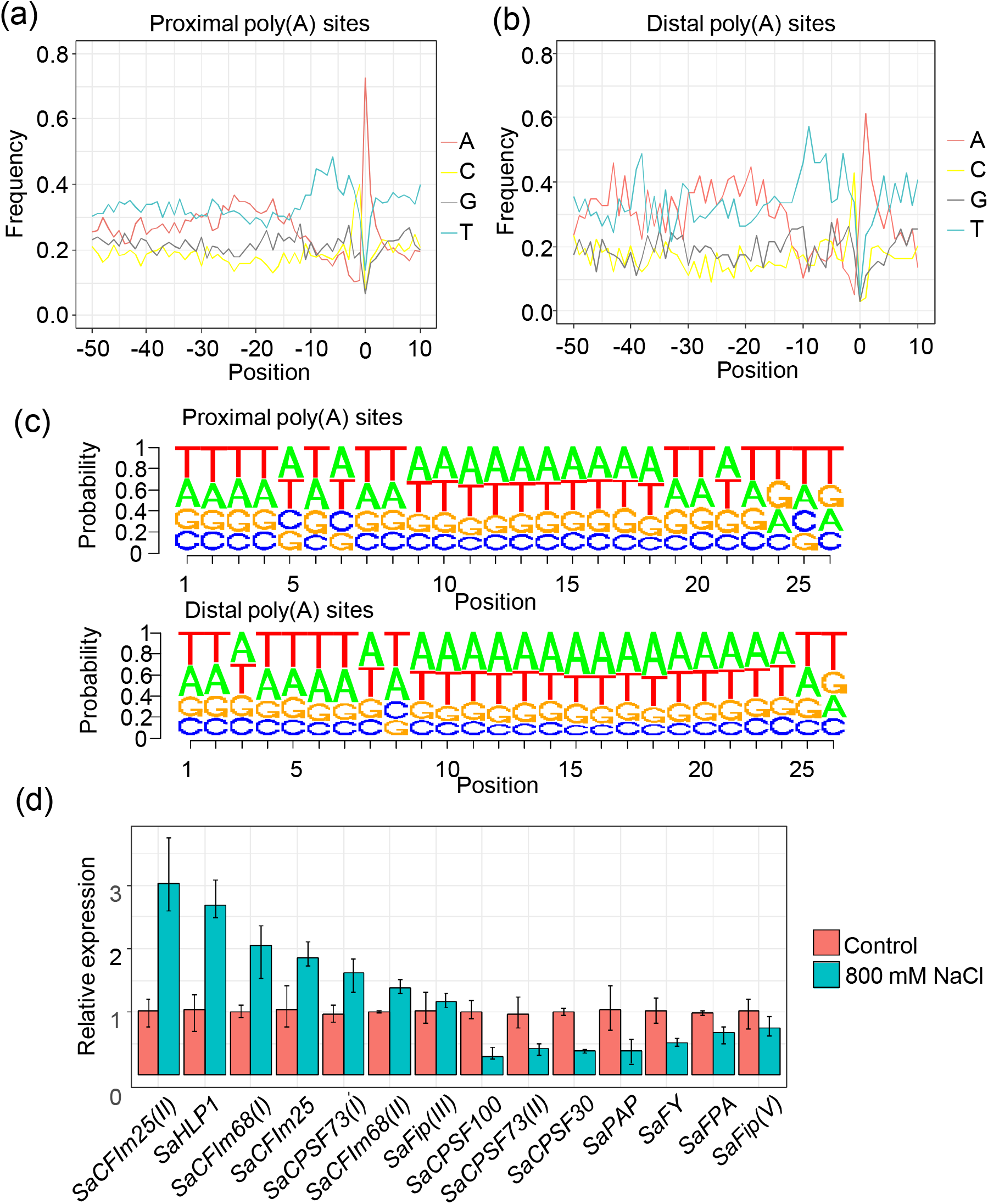
The biogenesis of 3′ UTR lengthening. (a) Nucleotide compositions of sequences surrounding proximal poly(A) sites of 3′ UTR lengthening transcripts. (b) As in (a) but for distal poly(A) sites. Y-axis denotes the fractional nucleotide content at each position. On the x-axis, “0” denotes the poly(A) site. (c) The nucleotide probability of sequences between –35 and –10 nt surrounding the proximal and distal poly(A) sites of the 3′ UTR lengthening transcripts. (d). The relative expression of *Spartina* poly(A) factors detected by qRT-PCR.

To test whether high salt stress affects the expression of poly(A) factors, gene expression levels of major poly(A) factors were determined by qRT-PCR under high salt stress. Importantly, 11 of the 14 detected genes showed an increase or decrease in gene expression under high salt stress, indicating that gene expression of poly(A) factors is regulated under high salt stress in a complex manner (Figure 6d). In particular, expression levels of two cleavage factors (*SaCFIm25* and *SaCFIm68*) increased, while *SaCPSF100* and *SaCPSF30* were down-regulated under high salt stress in *Spartina* (Figure 6d). Overall, the biogenesis of 3′ UTR lengthening may be caused by changes in poly(A) signal conservation and poly(A) factor expression under high salt stress.

## DISCUSSION

APA is one important post-transcriptional regulatory mechanism used by plants to respond to different environmental stresses (Deng and Cao, 2017; Sun et al., 2017; Srivastava et al., 2018). In this study, transcriptome-wide analysis indicated that APA involves in response to high salt stress (Figures 3-4). High salt stress induces a higher number of differential used APA sites along salt stress gradients (Figure 3a), and ∼42% of APA sites are associated with high salt stress, which is much higher than the prevalence of AS events (∼10%) (Ye et al., 2020). In addition, we found that pathways that are related to ion transport, response to osmotic stress, and respond to salt stress are significantly enriched in high salt stress-associated APA (Figure 3c-d). Overall, our results reveal widespread APA associated with high salt stress and the prevalence of post-transcriptional regulation in *Spartina* under high salt stress.

### The cause of 3′ UTR lengthening

One explanation of the preference of distal poly(A) sites under high salt stress is the differential expressed crucial poly(A) factors. The CFIm25 and CFIm68 have been found to promote the use of distal poly(A) sites in mammalian studies (Masamha et al., 2014; Gennarino et al., 2015). Similarly, the expression levels of *CFIm25* and *CFIm68* were increased under high salt stress (Figure 6d), which may result in a preference for distal poly(A) sites under high salt stress and thus promoting 3′ UTR lengthening of mRNAs. Moreover, mutation of CPSF100 and CPSF30 correlates with 3′ UTR lengthening by decreasing the usage of proximal poly(A) sites (Xia et al., 2014; Lin et al., 2017). Their reduced expression under high salt stress also leads to the preferential usage of distal poly(A) sites (Figure 6d). However, as mutant of *CPSF100* in *Drosophila* also promotes the read-through transcripts (Lin et al., 2017), we cannot exclude the possibility that high salt stress may lead to 3′ UTR lengthening via transcriptional read-through.

### The effect of 3′ UTR lengthening

In mammalian, alternative 3′ UTRs can regulate mRNA stability, mRNA localization, mRNA translation, and protein localization (Mayr, 2016, 2017). The AU-rich element and relevant RNA-binding-proteins (RBP) play vital roles in mediating mRNA stability and translation. In this study, we showed that high salt stress induced 3′ UTR lengthening of salt-responsive genes including ion transporter (Figure 4d). More importantly, AU-rich elements were specifically identified in alternative 3′ UTRs but not canonical 3′ UTR regions (Figure 5g). The AU-rich element was initially considered to enhance mRNA decay and also repress translation in animal (Shaw and Kamen, 1986; Kruys et al., 1989; Mayr, 2019). However, AU-rich element can also positively regulate mRNA stability and protein synthesis (Kontoyiannis et al., 1999). For example, the AU-rich element of tumor necrosis factor (TNF)-α in mouse permanently decreases protein expression in nonhematopoietic cells but transiently increases mRNA stability and translation in hematopoietic cell (Kontoyiannis et al., 1999). Similarly, the 3′ UTR lengthening of ion transporter-SaHKT1 increased their mRNA stability under high salt stress (Figures 5d-e), suggesting that alternative 3′ UTR may promote RNA stability. Take together with the observation that protein level of SaHKT1 was increased under high salt stress in an AU-rich dependent manner (Figures 5f, h), our results suggest that the inclusion of AU-rich element by alternative 3′ UTR stablizes RNA and increases protein synthesis of crucial salt-responsive genes under high salt stress in *Spartina*. Future work in other halophytes has to be done to clarify whether it is a *Spartina*-dependent or halophyte-specific event.

The AU-rich element inside alternative 3′ UTRs can act as an on and off switch to enable rapid and dynamic control of RNA stability and protein synthesis. It was found that the diverse function of same AU-rich element in different environment may largely dependent on the binding of RBPs as RBPs can bind to 3′ UTR and recruit effector proteins in animals (Mayr, 2017; Mayr, 2019). Currently, over 10 RBPs were identified to interact with AU-rich elements in animals (Mayr, 2019). TTP or KHSRP binds to AU-rich elements to destabilization mRNAs by anchoring exosome complex to them (Chen et al., 2001; Lykke-Andersen and Wagner, 2005). However, HuR stabilize AU-rich containing mRNA probably by blocking binding of the exosome to them (Chen et al., 2001). Therefore, we could also hypothesize that the AU-rich elements may also stable mRNA under high salt stress in an RBP associated pattern in plants. Future work will focus on identifing which RBPs bind to AU-rich elements inside alternative 3′ UTRs to decipher the underlaying molecular mechanism of RBPs and AU-rich elements module mediated RNA stability once the transformation system is ready for *Spartina alterniflora*.

However, 3′ UTR lengthening could contribute to RNA stability or protein synthesis regulation independent of AU-rich elements. For example, the U/A-rich regions surrounding APA sites (−15 nt to −2 nt) tend to form RNA secondary structures in Arabidopsis (Ding et al., 2014) and distal poly(A) sites of lengthened 3′ UTRs at the same region are higher U-rich than proximal poly(A) sites from canonical 3′ UTRs (Figure 6a-b). This indicates that alternative 3′ UTRs may promote RNA stability by affecting RNA secondary structures. Furthermore, U-rich elements interact with poly(A) tails to stabilize mRNAs in yeasts (Geisberg et al., 2014); the inclusion of two U-rich elements in transcripts with longer 3′ UTRs (Figure 5g) might affect RNA stability via this mechanism. In plants, it is also showed that dehydration stress induces 3′ UTR extension and acts as a long-non-coding RNAs to down-regulate downstream gene expression (Sun et al., 2017). In this study, non-coding RNA features of alternative 3′ UTR was not evaluated due to lack of neighbor-gene information from full-length transcriptome data. Therefore, we could not exclude that there is also such regulation in *Spartina*, which needs further evaluation once the genome sequence of *Spartina* is ready.

## CONCLUSIONS

Taken together, our results reveal a relationship between APA and high salt stress. The potential for 3′ UTR lengthening under high salt stress to affect protein expression and mRNA stability provides insights into how a salt-responsive mechanism may be regulated without affecting coding sequences. In addition, the identification of alternative 3′ UTRs of ion-transporters will provide useful information for future research on halophytes and engineering salt tolerant plants by genetic manipulation of poly(A) site choice and crucial *cis-elements*.

## MATERIALS AND METHODS

### Plant Materials and Treatment

*Spartina alterniflora* seeds collection and seedling growth conditions were exactly same as described in our parallel study using PacBio full-length transcriptome and RNA-seq (Ye et al., 2020). Briefly, six-week-old *Spartina* seedling were treated at different gradients of NaCl solution for 24 hours with three biological replicates to mimic the control (0), lower (350 mM), medium (500 mM) and higher salt stress (800 mM) conditions.

### PAT-seq library construction and sequencing

Total RNAs were extracted from *Spartina* seedlings after salt gradient treatment with commercial kit (Tiangen, Cat.DP441). Two microgram qualified total RNAs (RNA integrity > 8) were used to construct PAT-seq libraries using the method described in previous study (Wang et al., 2017). After quality evaluation with Agilent 2200 and determined concentration by qRT-PCR, the qualified libraries were pooled together to sequence on illumina HiSeq 2500 platform. The single-end poly(A) tag reads were generated for further bioinformatics analysis.

### Identification of differentially used APA sites

The detailed bioinformatics protocol for analysis of polyadenylation without reference genome was documented in **Methods S1**. Briefly, PAT-seq reads were mapped to transcripts identified from the PacBio full-length transcriptome data by STAR (v0.6.2) (Dobin et al., 2013) to obtain poly(A) sites. Internal priming artifacts were removed and poly(A) site clusters (PACs) were defined as described previously (Wu et al., 2011). PACs supported by at least 10 PAT-seq reads were retained for further analysis. Differentially expressed APA transcripts were identified using DEXseq (Anders et al., 2012) for clusters with at least two isoforms having 3′ reads. The expression level of the poly(A) site is the count of 3′ reads and expression levels were normalized between the respective conditions using the embedded function in DEXseq. For each pair of conditions (e.g., 350 mM vs. 800 mM), poly(A) sites with FDR<0.05 were identified as differentially used between the two conditions.

### Analysis of alternative 3′ UTRs

TransDecode (v3.0.1, https://transdecoder.github.io/) was employed to predict the 3′ UTR for each isoform. The distance from poly(A) site to stop codon was calculated as the 3′ UTR length for each poly(A) site. For each cluster, the read-weighted 3′ UTR length in a given condition (e.g., 350 mM) was calculated as the average 3′ UTR length of all isoforms weighted by the number of supported 3′ reads in this condition. Significant 3′ UTR lengthening or shortening between two conditions were detected by a test of linear trend (Fu et al., 2011; Fu et al., 2016). A correlation value (from −1 to 1) was calculated for each unigene to indicate the extent of 3′ UTR shortening (<1) or lengthening (>1). Unigenes with adjusted p-values smaller than 0.05 were considered as having significant 3′ UTR shortening or lengthening between two conditions, which were defined as APA-site switching unigenes.

### Analysis of poly(A) signals

To measure the relative base composition of poly(A) sites on a position-by-position basis, upstream 100 nt and downstream 100 nt sequence of each poly(A) site was extracted. Sequences of upstream 10 nt to 35 nt region of poly(A) sites were extracted to scan for the canonical poly(A) signal AAUAAA and its 1-nt variants. For comparison, poly(A) sites from other species were downloaded from the PlantAPA database (Wu et al., 2016).

### GO enrichment analysis

For each cluster, only the GO annotation of the longest isoform was used for GO enrichment analysis. The GO enrichment analysis for a given group isoforms was performed by BiNGO (Maere et al., 2005) using annotations of all longest isoforms as the background. GO terms with p-value less than 0.05 were considered to be enriched.

### RNA stability assay

The RNA stability experiment was performed according to the previous study in rice (Park et al., 2012). The *Spartina* seedlings were grown on half-strength Hoagland liquid medium for six weeks and then treated with 0 or 800 mM NaCl for 24hr. The *Spartina* seedlings were then grown in incubation buffer (1 mM PIPES, pH 6.25, 1 mM trisodium citrate, 1 mM KCl, 15 mM Sucrose) for 30 minutes. Cordycepin (3′-deoxyadenosine, Sigma, Cat.C3394) was used to treat the *Spartina* seedlings in a time course series (0, 30, 60, 90 and 120 minutes) analysis.

Total RNAs were then isolated (Tiangen, Cat.DP441). Steady-state mRNA levels were measured by qRT-PCR analysis using total RNAs from salt stress-treated *Spatina* seedlings. EIF-4A (Cluster1009-001) was used as an internal control to normalize the mRNA levels. Each means came from three independent qRT-PCR experiments and are presented relative to the results from unstressed controls with values set at 1. The changes of mRNA abundance were quantified over a time-course-experiment after exposure to 1 mM cordycepin. The total transcripts remaining was quantified using primers located on coding region and thus represented the abundance of transcripts with both the canonical and alternative 3′ UTRs (cUTR+aUTR); The primers located on aUTR regions were used to only quantified the remaining transcripts with alternative 3′ UTRs.

### Northern blot

The equivalence amount of total RNAs (15 μg) from six-week-old *Spartina* seedlings with or without high salt stress (800 mM) were used for RNA blot. The northern blot was performed as previous study (Knipple et al., 1998). A 288-bp digoxigenin (DIG)-labeled hybridization probe (alkali-labile digoxigenin-dUTP) was generated by PCR (35 cycles of 94°C for 30 s, 55°C for 30 s, and 72°C for 20 s) containing the *Spartina* cDNAs with the PCR DIG Probe Synthesis Kit (Roche, Cat.11636090910) using the *SaHKT1* transcripts specific primers (Forward primer: 5 ′-ACATCCTCACGAGACTGGCTAC-3 ′ and Reverse primer: 5 ′-AGCGAAGTATCAGAAGGAAGGT-3′). The RNA level was detected using the DIG-High Prime DNA Labeling and Detection Starter Kit II (Roche, Cat. 11585614910).

### Western Blot

Samples for protein expression analysis were exactly harvested and treated as samples from northern blot and RNA stability assay. The Anti-SaHKT1 polyclonal antibodies were raised in rabbits using the peptides C-KEENPEPAPSAPHQIQRVE as an antigen (ABclonal, Cat.WG-04220). The anti-Actin (Abbkine, Cat.A01050) was used as loading control. Total proteins were extracted by the Minute™ Total Protein Extraction Kit for Plant Tissues (Invent, Cat.SN-009). 40 μg total proteins were loaded into 10% acrylamide gels and transferred to Immobilon-P membrane (Millipore, Cat. IPVH00010). The antibody were diluted as follows: Anti-SaHKT1(1:1000) and anti-Actin (1:3000). The protein levels were detected by using SuperSignal^TM^ West Pico Plus (ThermoFisher, Cat.34577) following manufacture’s recommendation.

### Protoplast Isolation and Dual-luciferase assay

The *Spartina* protoplast transient expression protocol was modified from previous study in moso bamboo (Lin et al., 2021). *Spartina* seeds preserved in 1.5% sea salt water were washed with double distilled water and growth on the filter papers inside petri dishes on a dark condition at room temperature (25℃) for one week to reach 10 cm tall. After removing the root, 200 fresh and tender yellow seedlings were collected and cut into 1-2 mm fragments.

The longer and shorter transporter 3′ UTRs were cloned into the dual luciferase construct pGreen_dualluc_3′UTR_sensor at *Eco*RI sites (Addgene, Cat: 55206). The longer 3′ UTRs without AU-rich elements were amplified using flanking PCR methods. The constructs were transformed into *Spartina* protoplasts in an PEG-Calcium-dependent transfection. A dual luciferase assay was performed in *Spartina* protoplasts using the Dual Luciferase Reporter Assay system (Promega, Cat: E1980), as described previously (Liu et al., 2014). Firefly and Renilla luciferase were measured using a microplate luminometer (Berthold Technologies, Centro XS LB960). The detailed procedures of protoplast transformation and dual-luciferase assay were documented in **Methods S1**.

### Experimental validation

The bench experiments including 3′ RACE and qRT-PCR experiments were conducted as described in previous study (Wang et al., 2017) and the results were visualized in 1% agarose gels. Primers used in this study are listed in **Table S4**. Full methods in this study are documented in **Methods S1.**

### Data availability and accession numbers

Datasets from Illumina HiSeq 2500 sequencing for PAT-seq have been deposited at the NCBI website under the Bioproject accession number PRJNA413596. Alternative polyadenylation can be visualized in our SAPacBio website (http://plantpolya.org/SAPacBio/).

## Competing financial interesting

The authors declare no competing financial interests.

## Acknowledgements

This work was supported by the National Natural Science Foundation of China (31741025 and 31500258 to L.M., and 61673323 to X.W.), the Natural Science Foundation of Fujian Province of China (2020J01592 to L.M.), the Special Fund for Science and Technology Innovation of Fujian Agriculture and Forestry University (CXZX2019142G to L.M.).

## Author Contributions

X.W. and L.M. conceived the original research plans; G.J. supervised the experiments; T.W. performed the sample preparation, Northern blot, 3′ RACE, qRT-PCR, RNA stability assay and molecular cloning; W.Y. performed all bioinformatics analyses; J.Z. established the *Spartina* protoplast transient expressed system and performed dual-luciferase assay; H.L. performed Western blot analysis; W.Z. performed all statistical analyses; S.Z. established and maintained the *Spartina* website; L.M. wrote the article. All authors read and approved the final manuscript.

## SUPPORTING INFORMATION

### Supplemental Figures

**Figure S1.** Distribution of distances from PACs located in the 3′ UTR to the 3′ end of respective full-length transcripts.

**Figure S2.** Number of genes with alternative 3′ UTRs. Each bar denotes the number of genes with longer or shorter 3′ UTRs in the respective sample.

**Figure S3.** Relative expression of salt-responsive unigenes under high salt stress detected by qRT-PCR.

### Supplemental Tables

**Table S1.** Summary of statistical analysis of PAT-seq data.

**Table S2.** List of differentially expressed PACs for transcripts with multiple poly(A) sites.

**Table S3.** List of unigenes with significant 3′ UTR lengthening under high salt stress.

**Table S4.** List of primers used in this study.

**Supplemental Methods:** Full methods were documented in Methods S1. Figure legends

