## Supplemental Figures for "Alternative 3′ UTRs contributes to post-transcriptional gene expression regulation under high salt stress"

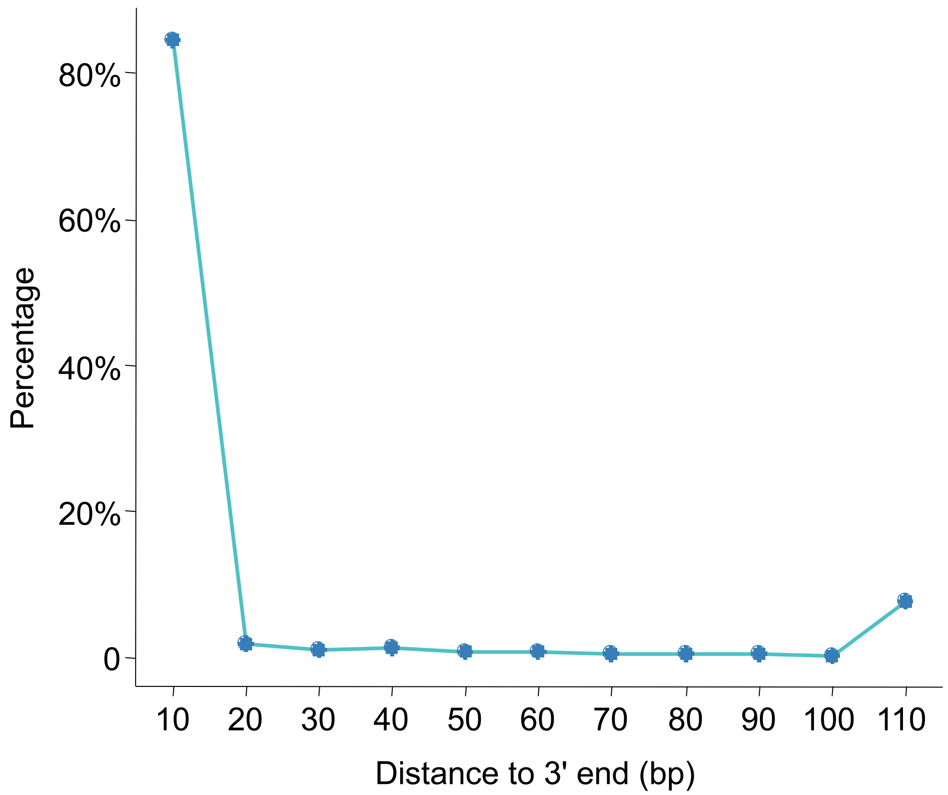


Figure S1 Distribution of distances from PACs located in 3′ UTR to the 3′ end of respective full-length transcripts.

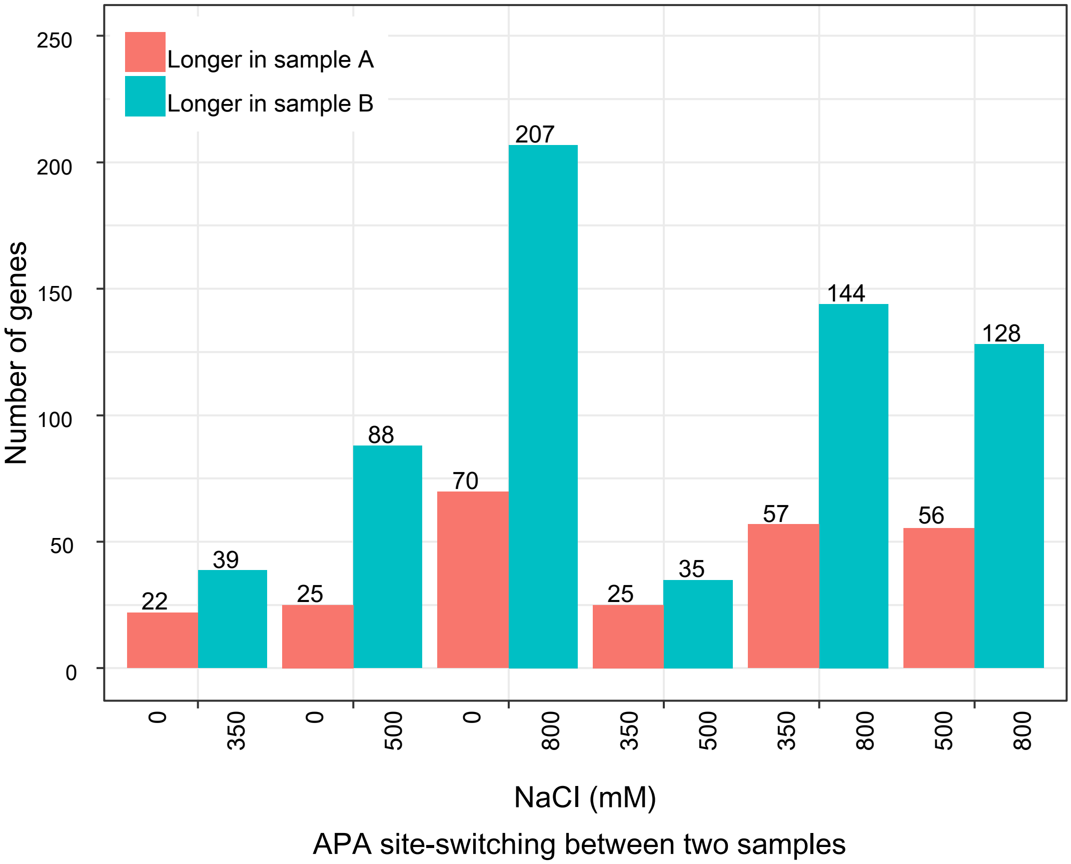


Figure S2 Number of genes with alternative 3′ UTRs. Each bar denotes the number of genes with longer or shorter 3′ UTRs in the respective sample.

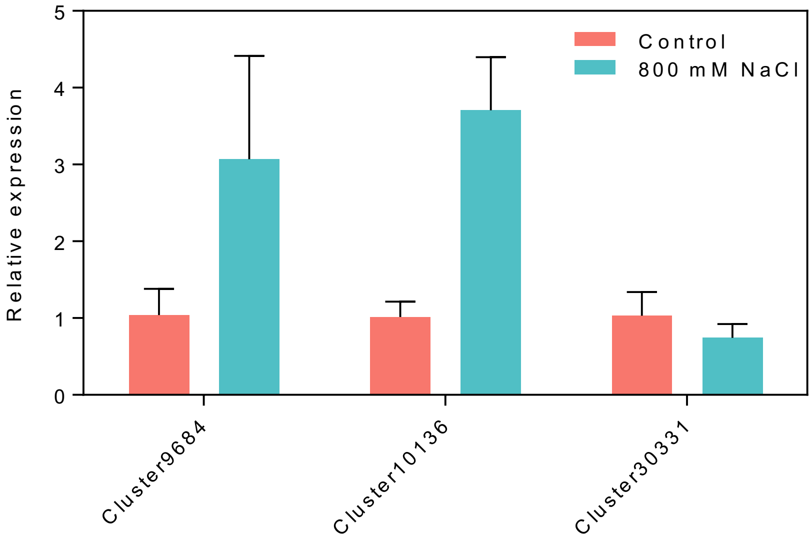


Figure S3 Relative expression of salt-responsive unigenes under high salt stress detected by qRT-PCR.
