## Supplemental Methods for "Alternative 3′ UTRs contributes to post-transcriptional gene expression regulation under high salt stress"

**Analysis of APA without a reference genome**

**Pre-processing of PAS-seq data**

The PAS-seq data were sequenced from two batches with different read length (30 bp and 50 bp). To make the results consistent, longer reads were trimmed by 20 bp from the 3' end for subsequent analysis. The stretches of Ts at the beginning of the reads were trimmed by a Perl script. Trimmed reads with at least 25 bp were retained and subsequently aligned to full-length isoforms using STAR (v0.6.2, parameters: --outMultimapperOrder Random --outSAMmultNmax -1 --outFilterMultimapNmax 200 --outFilterMismatchNmax 10) . Files in SAM format were generated. It should be noted that using the longest isoform in each cluster as reference may neglect reads mapped to regions not present in the longest isoform. Therefore, in the mapping procedure, reads were mapped to all full-length isoforms detected from SMRT data and each read was allowed to match a maximum 200 positions. Reads that only mapped to one isoform cluster were retained and those that mapped to multiple isoform clusters were discarded. Reads that mapped to multiple positions of the same isoform were also removed. Additional filtering criteria were applied to filter effective reads for the respective analysis. SAM files were parsed and coordinates of poly(A) sites were obtained for each isoform. False poly(A) sites of internal priming were removed by scanning the enrichment of A around the mapped position based on a previous protocol .

**Determination of distal poly(A) sites for full-length isoforms**

For a read that mapped to multiple isoforms in one cluster, we determined the most probable isoform that it originated from, based on the distance from the mapped position to the 3' end for each isoform (Methods Fig. **1**). If the difference of the distances to the 3' end of two isoforms of a read was less than 10 bp and, only if one isoform was the longest isoform in this cluster (the unigene defined in this study), then the longest isoform was assigned to this read, otherwise the read was discarded. If the distance for mapped isoforms of a read was distinguishable (> 10 bp), given that a poly(A) site is more likely to be present at the 3' end of an isoform than in the body region, the read was assigned to the isoform with the shortest distance. The 3' poly(A) reads of an isoform were defined as reads located within the upstream 100 bp region of the isoform's 3' end. The number of 3' reads for each isoform was used as the expression level for further analysis.

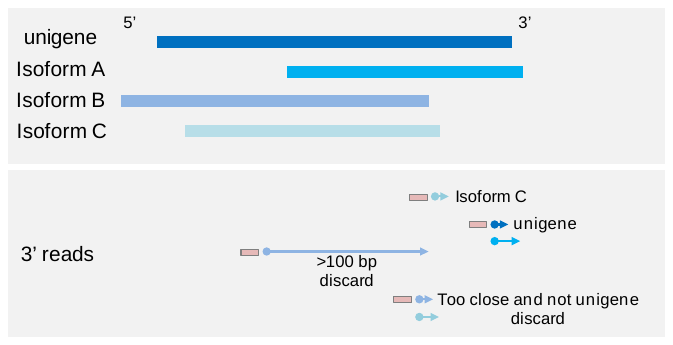

**Methods Fig. 1** Schematic diagram of determining 3' end poly(A) sites for each full-length isoform in a cluster.

**Identification of differentially used APA sites**

Because no reference genome is available and using the longest isoform to represent each cluster would neglect poly(A) sites not present in the longest isoform, we obtained the distal poly(A) site for each isoform in a cluster if the isoform was supported by at least ten 3' reads. DEXseq , which was originally developed to detect differential exon usage from RNA-seq data, was employed to identify differentially used APA sites. DEXseq has been proven to be effective in identifying signiﬁcant changes in 3' UTR isoform expression . To use DEXseq, only clusters with at least two isoforms having 3' reads were used. Each cluster was considered as a gene and the 3' end poly(A) sites for all isoforms with 3' reads were considered as exons. The expression level of the poly(A) site is the count of 3' reads and expression levels were normalized between the respective conditions using the embedded function in DEXseq. For each pair of conditions (e.g., 350 mM vs. 800 mM), poly(A) sites with FDR<0.05 were identified as differentially used between the two conditions.

**Analysis of 3' UTR lengthening and shortening and identification of APA site switching unigenes**

TransDecode (v3.0.1, https://transdecoder.github.io/) was employed to predict the ORF, 5' UTR, and 3' UTR for each isoform (command: TransDecoder.LongOrfs -m 100 –S; TransDecoder.Predict --single_best_orf). Out of the 90,587 full-length isoforms, 84,715 (93%) carried complete ORF (with start and stop codons) over 100 amino acids. For each cluster, the read-weighted 3' UTR length in a given condition (e.g., 350 mM) was calculated as the average 3' UTR length of all isoforms weighted by the number of supported 3' reads in this condition. Isoforms without a 3' UTR predicted by TransDecode were excluded.

To discover APA site switching isoform clusters with significant 3' UTR shortening or lengthening across different conditions, only the longest isoform in each cluster was considered. Pre-processed PAS-seq reads that mapped to the longest isoforms were retained and coordinates of cleavage sites were obtained. Nearby cleavage sites from all the four conditions within 24 bp of each other were grouped as a single poly(A) site to eliminate the effect of micro-heterogeneity. Poly(A) sites supported by at least ten reads were retained and the number of reads in each experiment was recorded. Only poly(A) sites located in predicted 3' UTRs by TransDecode were used and isoforms with at least two poly(A) sites in the 3' UTRs were considered. The distance from poly(A) site to stop codon was calculated as the 3' UTR length for each poly(A) site. Significant 3’ UTR lengthening or shortening between two conditions were detected by a test of linear trend . A correlation value (from -1 to 1) was calculated for each unigene to indicate the extent of 3' UTR shortening (<1) or lengthening (>1). Unigenes with adjusted p-values smaller than 0.05 were considered as having significant 3' UTR shortening or lengthening between two conditions, which were defined as APA-site switching unigenes.

***Spartina* protoplast Protocol**

**The *Spartina* protoplast protocol was modified based on previous protocols.**

**MATERIALS**

**Reagents**

0.8 M mannitol (Solarbio, cat. no. M8140)

0.2 M 4-morpholineethanesulfonic acid (MES, PH=5.7) (Bomei, cat. no. MM8655)

1 M CaCl_2_ (XILONG, cat. no. 10301301)

2 M KCl (Sigma. cat. no. P9514)

2 M MgCl_2_ (Rhawn, cat. no. R007504)

10% (wt/vol) BSA (Solarbio, cat. no. A8010)

Cellulase R10 (Yakult Pharmaceutical Ind. Co., Ltd., Japan)

Macerozyme R10 (Yakult Pharmaceutical Ind. Co., Ltd., Japan)

Cellulase “Onozuka” RS (Yakult Pharmaceutical Ind. Co., Ltd., Japan)

PectolyaseY-23 (Yakult Pharmaceutical Ind. Co., Ltd., Japan)

PEG4000 (Macklin, cat. no. P815608)

LUC assay system (Vazyme, cat. no. DL101-01)

Enzyme solution (see REAGENT SETUP)

Washing and incubation (WI) solution (see REAGENT SETUP)

W5 solution (see REAGENT SETUP)

MMG solution (see REAGENT SETUP)

PEG–calcium transfection solution (see REAGENT SETUP)

Protoplast lysis buffer (Vazyme, cat. no. DL101-01)

**QUIPMENT**

Fluorescence microscope (Zeiss, Axio Observer.A1)

Cytation Hybrid Multi-Mode Reader (Bioteck, cytation5)

Bench-top centrifuge (Eppendorf, cat. no. 5810R)

0.45-mm syringe sterilization filter (Whatman, cat. no. 6870-2504)

Surgical blade (single-edged blade; VWR Scientific, cat. no. 55411-055)

Petri dish (Biosharp, cat. no. BS-90-D)

Nylon mesh (75 mm, laboratory sifters, Carolina Biological Supplies,cat. no. 65-2222N)

0.1-mm-deep Reichert hemacytometer (CHANGDE BKMAM BIOTECHNOLOGY CO.,LTD, cat. no. 1103)

50-ml round-bottomed tube (Corning, cat. no. 430828)

2-ml round-bottomed natural microcentrifuge tube (Biosharp, cat. no. BS-20-M)

6-well culture dish (Biofil, cat. no. 3046)

96-well micrp plate(corining)

Inoculation loop (BD, cat. no.220217)

filter paper (GE, cat. no. 3030704)

**REAGENT SETUP**

**Enzyme solution**

Prepare 20 mM MES (PH5.7) containing 1.5% (wt/vol) Cellulase R10, 0.2% (wt/vol) Macerozyme R10, 0.2% (wt/vol) Cellulase “Onozuka” RS, 0.2% (wt/vol) PectolyaseY-23, and 0.4 M mannitol. Mix well and heat in water bath at 55ºC for 10 min and wait for room temperature to add 10 mM CaCl_2_. Use a 45 μm filter to filter and sterilize.

**WI solution**

Prepare 4 mM MES (PH = 5.7) containing 0.5 M mannitol and 20 mM KCL. The prepared WI solution is autoclaved and stored in a refrigerator at 4°C.

**W5 solution**

Prepare 2 mM MES (PH = 5.7) containing 154 nM NaCl, 125 nM CaCl_2_ and 5 nM KCl, The prepared W5 solution is autoclaved and stored in a refrigerator at 4°C.

**MMG solution**

Prepare 4 mM MES (PH = 5.7) containing 0.4 M mannitol and 15 mM MgCl_2_. The prepared W5 solution is autoclaved and stored in a refrigerator at 4°C.

**PEG–calcium transfection solution**

Prepare 20%-40%(wt/vol) PEG4000 in ddH_2_O containing 0.8 M mannitol and 15 mM MgCl_2_.

**PROCEDURE**

**Plant growth _ TIMING 1 week**

*Spartina alterniflora* seeds preserved in 1.5% sea salt water are washed with double distilled water. Placed the seeds on filter paper to a 9*9 square petri dish, and then irrigated with double distilled water to avoid light at room temperature (25°C) for a week. During this period, water the seedlings continuously until the stems and leaves of *Spartina alterniflora* grow to 10 cm tall.

**Protoplast isolation**

(1) Choose 200 fresh and tender yellow seedlings from 1 week, cut remove the root and cut 1-2 mm of the whole seeding by using a fresh surgical blade without crushing tissue at the cut site.
(2) Gently transfer the stem and leaf fragments to the round petri dish with the enzymatic hydrolysate. And gently tap the stems and leaves to submerge them.
(3) Use a vacuum dryer to infiltrate the stems and leaves in the dark for 30 minutes.
(4) Continue enzymatic hydrolysis, wrap the Petri dish with tin foil and place it on a horizontal shaking table with a rotating speed of 50 rpm.
(5) Check cell morphology and size under a microscope.
(6) According to the volume of the enzymatic hydrolysis solution, add an equal volume of pre-cooled W5 solution to terminate the enzymatic hydrolysis reaction.
(7) Wash the 75 nm nylon mesh with pure water to remove the absolute ethanol. Try to remove the excess water before filtering.
(8) Centrifuge horizontally at 1300 rpm (Acceleration and braking acceleration is 3) for two minutes in a 50 ml round bottom centrifuge tube. Try to remove the supernatant, add 10 ml of pre-chilled W5 solution and slowly add it along the tube wall. Centrifuge horizontally at 1300rpm again.
(9) Add 10 ml of pre-cooled W5 solution slowly along the wall of the tube, Keep the protoplasts on ice for 30 minutes.
(10) Centrifuge at 1300 rpm for 2 min to remove the supernatant solution as much as possible. Add the required volume of MMG solution to resuspend and keep the protoplasts. Observe and count protoplasts under a microscope.

**DNA-PEG–calcium transfection**

(11) Add 200 μl protoplasts to a 2-ml microfuge tube，and then add 20 μl DNA (1000 ng/μl of plasmid DNA) to each tube and mix gently.
(12) Add 220 μl of PEG solution and mix gently and thoroughly. Incubate the mixture for 5 min at room temperature. Gently add 1 ml of W5 solution to the transfection mixture at room temperature. Mix gently to stop the conversion reaction.
(13) Centrifuge at 1300 rpm for 2 min to remove the supernatant.
(14) Add 100 μl of W5 solution to resuspend the protoplasts, and transfer to a six-well dish rinsed with 10% BSA and filled with 1 ml WI solution.

**Protoplast culture and harvest**

(15) Incubate the protoplasts at room temperature (20-25℃) for 12 hours while giving continuous light.
(16) Centrifuge at 1300 rpm for 2 min to remove the supernatant solution and resuspend protoplasts

**Dual-luciferase assay**

(17) Add 45 μl protoplast lysis buffer for each tube and mix to repute the protoplasts for 5 min.
(18) After 5 incubation centrifuges at 12000 rpm for 2 min, then divide into 96-well plates as needed.
(19) Add firefly luciferase and renilla luciferase, which are equilibrated to room temperature, into the 96-well microplate, and measure the fluorescence intensity in the microplate reader.
